# Flaw or discovery? Calculating exact p-values for genome-wide association studies in inbred populations

**DOI:** 10.1101/015339

**Authors:** Xia Shen

## Abstract

**Motivation:** Genome-wide association studies have been conducted in inbred populations where the sample size is small. The ordinary association p-values and multiple testing correction therefore become questionable, as the detected genetic effect may or may not be due to chance, depending on the minor allele frequency distribution across the genome. Instead of permutation testing, marker-specific false positive rate can be analytically calculated in inbred populations without heterozygotes.

**Results:** Solutions of exact p-values for genome-wide association studies in inbred populations were derived and implemented. An example is presented to illustrate that the marker-specific experiment-wise p-value varies as the genome-wide minor allele frequency distribution changes. A simulation using real *Arabidopsis thaliana* genome indicates that the use of exact p-values improves detection power and reduces inflation due to population structure. An analysis of a defense-related case-control phenotype using the exact p-values revealed the causal locus, where markers with higher MAFs had smaller p-values than the top variants with lower MAFs in ordinary genome-wide association analysis.

**Availability and Implementation:** Project URL: https://r-forge.r-project.org/projects/statomics/. The R package **p.exact:** https://r-forge.r-project.org/R/?group_id=2030.

## 1 INTRODUCTION

As high-throughput genotyping technologies are currently available in many species, genome-wide association studies (GWAS) have been conducted in various kinds of populations for the purpose of discovering genomic loci associated with complex traits. One particular type of population consists inbred individuals, which are either due to natural selfing (in plants) or artificially created inbred lines. Such inbred populations are genetically powerful resources in association analyses as no heterozygotes exist, so that the genotype data are less heterogeneous, thus noise is reduced. However, obtaining an inbred individual often requires more effort than simply collecting a large number of outbred individuals. Therefore, GWAS conducted in inbred populations are usually based on a small number of individuals, but the number of single nucleotide polymorphism (SNP) markers tested is large.

This brings up a challenge in terms of reporting GWA signals with sufficient confidence. For instance, if only 20 inbred lines are studied, and a significant association is found at a SNP where 5 lines with genotype *AA* have the largest 5 phenotypic values and the other 15 *CC* have the rest, the genetic effect seems rather clear. But because plenty of markers are tested across the genome, e.g. 100 000, the chance of a random marker splitting the 20 phenotypic values into such two groups does not feel low. One can perform a permutation test to overcome such uncertainty, but unfortunately due to computer intensity, a Bonferroni-corrected significance threshold is still often applied, which is regarded as “conservative” in literature. On the other hand, discoveries based on low MAFs may not be as questionable as we normally think. Simulation studies have indicated that in certain genomes, the false discovery rates of rare variants may not be as high as expected (Tabangin *et al*., 2009).

In fact, as only two homozygotes exist in an inbred population, the number of observed minor allele frequencies (MAFs) across the genome is rather small, so that the exact count of each MAF value can be obtained. Using this genome-wide MAF distribution, the exact marker-specific false positive rate (p-value) has an analytical solution, which can substantially save the computational time needed in a permutation test. The aim of this paper is to derive and implement this analytical solution of exact p-values and emphasize the use of such a tool in inbred-population GWA analyses to prevent future studies from reporting flawed results.

## 2 METHODS

For *n* inbred individuals, consider a phenotypic vector y on a continuous scale to be studied, adjusted for confounding covariates and inverse-Gaussian transformed, and *k* SNPs across the genome. There are only

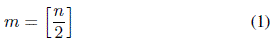

different values of observed MAFs. For a particular SNP with major allele *Q*, minor allele *q*, and MAF = *f*, there are *n f* individuals having the genotype *qq* and *n*(1 – *f*) individuals having *QQ* at this marker. The observed (estimated) absolute effect size is

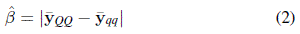

When *n* or *f* is small, the estimated standard error of 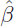 is not reliable in such high-throughput data analysis with severe multiplicity. Instead, we can infer the p-value for such an effect size based on the sample size *n* and the MAF distribution across the genome.

It can be counted that there are *k*_1_ SNPs with MAF *f*_1_, *k*_2_ SNPs with MAF *f*_2_, …, *k_m_* SNPs with MAF *f_m_*, and 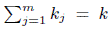. Now consider each of the *k_j_* SNPs with MAF *f_j_*, there are *nf_j_* individuals having the minor genotype and the other *n*(1 – *f_j_*) having the major. Under the null, i.e. no SNP is associated with the phenotype, we have

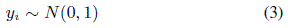

for each individual *i*, where *N*(*μ, σ*^2^) denotes a Gaussian distribution with mean *μ*, and variance *σ*^2^. Therefore,

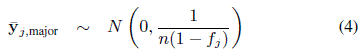

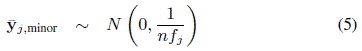

and

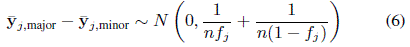

We have that the probability of observing an absolute SNP effect less than |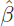| for a SNP with MAF *f_j_* is

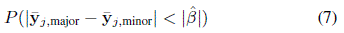

for the same effect size, ∀*j*\* *s.t. f_j*_* < *f_j_*, we have

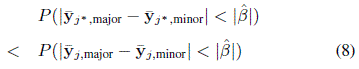

Thus, given 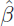, false discoveries can only be generated at a SNP with MAF ⩾ *f_j_*. Using the observed MAF distribution,

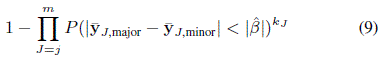

is the exact dataset-specific p-value for 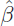. The p-value from equation (9) is two-sided, and the one-sided option is also available in the software package.

Clearly, direct implementation using equation (9) involves multiplication of a large number of values between 0 and 1, and therefore suffers from numerical difficulty. Alternatively, the p-value can be calculated as

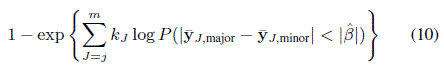

Also, it should be noted that such a p-value has already accounted for multiplicity in the genome-wide association study. Namely, the p-value of each SNP indicates the false positive rate for the whole experiment, which is the analytical exact solution of a computer-intensive permutation test by re-shuffling the phenotypic vector.

If y is a binary vector, e.g. in case-control studies, we have

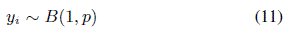

under the null, and the odds ratio (OR*_s_*) between the major and minor alleles of a particular SNP *s* has a mean of 1. Thus, if the corresponding 2 × 2 contingency table is

|  | Major | Minor |
| --- | --- | --- |
| Case | $n_{s11}$ | $n_{s12}$ |
| Control | $n_{s21}$ | $n_{s22}$ |

we have

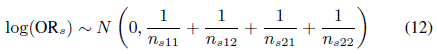

Similarly, given that *f_s_* is the MAF of SNP *s*,

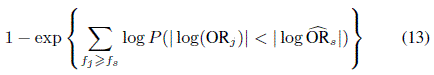

is the exact p-value for 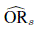. When zero count exists in the contingency table, replacing 0 with 0.5 keeps the validity of the asymptotic distribution (12).

The above calculations for continuous and binary phenotypes are implemented as two separate modules in the **p.exact** package on R-Forge. The package incorporates the `gwaa.data` class objects of the **GenABEL** package (Aulchenko *et al*., 2007) which is commonly used in R-based GWA analyses, compatible with different types of data formatting.

## 3 EXAMPLES AND RESULTS

Simulations were used to illustrate the performance of the exact p-values. In an inbred population with sample size *n* = 100 and the number of SNPs *k* = 100000, a Bonferroni-corrected 5% genome-wide significance threshold is 5.0 × 10^−7^. For a SNP with MAF = 0.05 and an estimated effect size 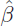 = 2 on a normalized phenotypic scale, the ordinary GWA p-value is

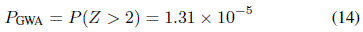

where *Z* ~ *N*(0, 1/5 + 1/95). Thus, such a marker would not be reported as significant in ordinary GWAS as long as *k* does not vary. However, the exact false positive rate (p-value) of this marker would depend on the MAF distribution across the genome, as shown in Figure 1. The more low-MAF markers exist, the more likely the observed effect is due to chance, and a higher p-value should be assigned to the SNP. Therefore, for a genome with MAF distribution as in Figure 1 (A, B, E, F), this marker has actually a significant experiment-wise exact p-value that is less than 0.05.

**Fig. 1.**
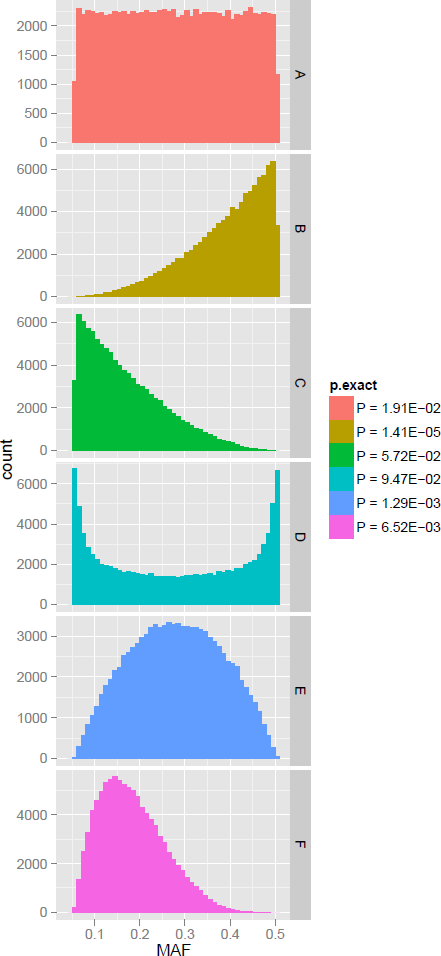
Dependency of the exact p-value on whole-genome allele frequency distribution. An inbred population with 100 individuals were simulated with 100,000 SNPs across the genome. Six different scenarios of minor allele frequency (MAF) distribution across the genome were simulated (A–F), where for each scenario, a SNP with MAF = 0.05 and estimated effect size 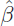 = 2 is tested using **p.exact**. From A to F, the ordinary GWA p-value is the same, i.e. 1.31 × 10^−5^, but the exact p-value based on the whole-genome MAF distribution differs.

Another simulation was conducted using a real *Arabidopsis thaliana* dataset (Atwell *et al*., 2010). In the SNP array, 216 130 markers were genotyped in 199 inbred accessions. Phenotypic values were simulated by randomly selecting a SNP as the single causal variant, with 0.05 narrow sense heritability (*h*^2^). Both a Gaussian and a binary (case-control) trait were simulated, and the process was repeated for 500 times to compare the p-values inferred by the GWA package **GenABEL** and **p.exact** (Figure 2). Mixed modeling for controlling population stratification was only used in analyzing the normally distributed trait. The simulation indicated that **p.exact** is more conservative when population stratification exists (Figure 2A) but also more powerful (less conservative) when the population structure is corrected (Figure 2B).

**Fig. 2.**
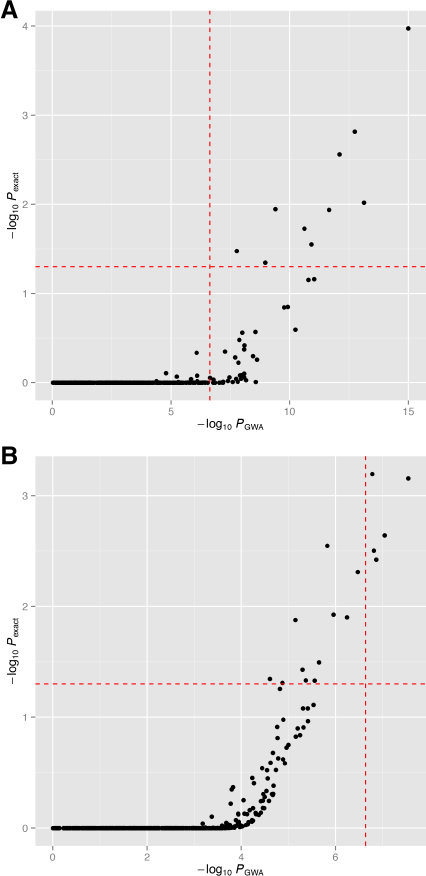
Comparison of the ordinary GWA and exact p-values for a simulated QTL. The genomic SNP data from Atwell *et al*. (2010) containing 199 accessions and 216 130 SNPs were used. A SNP was randomly selected from the genome to be the causal variant of the QTL, with narrow sense heritability = 0.05. Such simulation procedure was repeated for 500 times. The vertical and horizontal dashed lines represent the Bonferroni corrected 5% significance threshold in ordinary GWAS and 5% threshold for the exact p-values, respectively. (A) A binary (case-control) trait was simulated in each replicate, and *P*_gwa_ was obtained via the `ccfast` procedure in **GenABEL;** (B) A normally distributed trait was simulated, and *P*_gwa_ was obtained using the `qtscore` procedure in **GenABEL**, with GRAMMAR+ (`grresidualy` from `polygenic`; Belonogova *et al*., 2013) transformed phenotype to control for population stratification.

For a further comparison, in the same *Arabidopsis thaliana* dataset, a real case-control disease-resistance trait AvrRpm1 was analyzed using both ordinary GWAS and the exact p-values for all the markers across the genome (Figure 3). The single causal locus *RPM1* was mapped using both methods, however, different sets of markers were above the significance thresholds. If based on single marker nominal p-values, the variants with MAF = 0.25 in the LD (linkage disequilibrium) block had the smallest p-values. While considering the whole-genome MAF distribution, the exact p-values highlight the variants with more common minor allele (MAF = 0.37) instead, as our confidence on the signal drops if the finding is based on small MAF values.

**Fig. 3.**
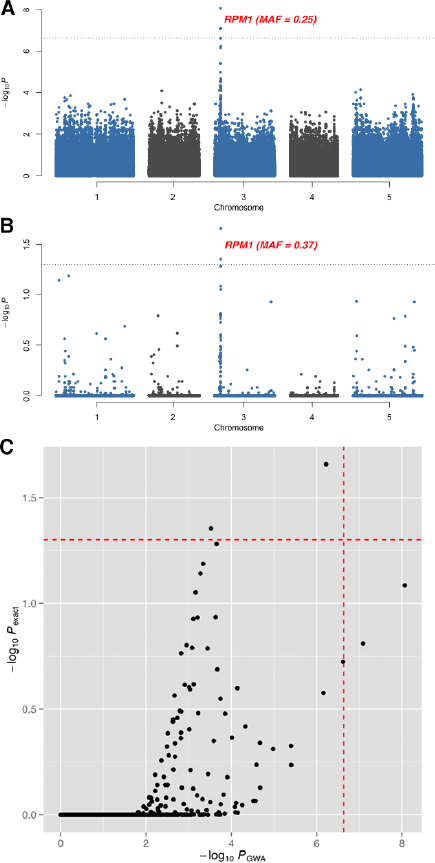
Comparison of (A) the ordinary GWA and (B) exact p-value scans of the *Arabidopsis* bacteria-hypersensitive trait AvrRpm1 (Atwell *et al*., 2010), with 84 accessions and 215 130 SNP markers. *P*_gwa_ and *P*_exact_ were obtained by the `ccfast` procedure in **GenABEL** and `p.exact.binary` procedure in **p.exact**, respectively. The causal locus *RPM1* is labeled in (A) and (B), with the MAF of the top variants. The vertical and horizontal dashed lines in (C) represent the Bonferroni corrected 5% significance threshold in ordinary GWAS and 5% threshold for the exact p-values, respectively.

## 4 DISCUSSION

Described here is an exact false discovery rate calculation, accounting for the overall allele frequency property in each high-throughput GWA dataset. Although the mathematics behind the calculation is relatively “trivial” and limited to inbred populations, it provides an *analytical* solution of an equivalent computer-intensive permutation test, which is often intractable in general outbred populations. Besides efficiency, its usage should be emphasized in small sample association studies, which are mostly conducted in small inbred populations, where “discoveries” are likely flawed due to multiple testing a small number of individuals.

The presented tool can also be useful when screening the genome for epistasis, i.e. gene × gene interactions, where multiplicity is more problematic. In that case, the frequency of the minor-minor genotype combination is crucial, so that **p.exact** can be used to test the corresponding two-way “pseudo markers”. In one of our applied work (Lachowiec *et al*., 2014), we were able to detect four pairs of epistasis with highly significant (using nominal p-values) effects, however, the exact p-values described here statistically declared three pairs to be flawed discoveries, simply because too many “pseudo markers” had minor-minor genotype frequency as low as the three pairs (0.043). Such a problem may not be vital in large sample GWAS, especially in outbred human cohorts, but could be severely misleading in small populations.

Extension of the exact p-value calculation to outbred populations can be possible if strict Hardy-Weinberg equilibrium is assumed for each SNP. But such an assumption is too strict for real data. Also, unlike the inbred samples, there are not many markers with exactly the same MAF across the genome, namely, the number of possible MAF values across the genome in an inbred population is very limited. Therefore, it is in inbred populations that the analytical solution here is well defined, and it is also in inbred (often small) populations that the risk of reporting flawed results is considerably high.

Although the analytical solution is exact, the method can still be conservative due to high LD with each locus. Nevertheless, in the implemented **p.exact** package, an LD-pruning option is available for excluding adjacent high LD markers. When testing this procedure on the *A. thaliana* genome, the results did not differ much, as in the current 250K *A. thaliana* SNP array, there are not so many high LD markers within each locus. Only 6% of the markers can be excluded with an LD *r*^2^ cutoff of 0.90. Nevertheless, high LD-pruning will be essential if more dense markers or whole-genome sequencing variants are analyzed.

## ACKNOWLEDGEMENT

The author thanks Chris S. Haley for helpful discussions.

### Funding

This work is supported by a Swedish Research Council grant (537-2014-371) to XS.

